# Mite Genome Miniaturization: Assembly of the biological control agent *Floracarus perrepae* reveals dynamic genome evolution in eriophyoid mites

**DOI:** 10.64898/2026.08.24.746826

**Authors:** Jessie A. Pelosi, Taylor R. Curry, Melissa C. Smith, Katrina M. Dlugosch

## Abstract

The eriophyoid mites (Acari: Eriophyoidea) represent an extreme case of genome streamlining, with genomes averaging just 32Mb, among the smallest of all animals. This clade of mites is highly diverse with more than 4000 species. Their morphologies are specialized for feeding on their host plants, including a simplified worm-like body plan with just two pairs of legs and modified mouth parts that can induce the formation of galls or plant deformities during feeding. Their generally strong host affinities make eriophyoid mites appealing for use as biological control agents, although their short generation times and small genomes could facilitate rapid evolution and impact their efficacy in management programs. Here, we sequenced the genome of the biological control mite *Floracarus perrepae*, producing a highly contiguous genome totalling just 23.1 Mb, among the smallest of all animals. We also assembled another non-eriophyid mite genome from accidental DNA bycatch (69.4Mb). We placed this new genomic resource in a phylogenetic context to reveal that mites have highly dynamic genome evolution, with a significant trend in genome downsizing in the eriophyoids. Our results suggest that this streamlining is associated with non-genic elements such as the suppression or excision of retrotransposons and purging of introns. As new sequencing techniques become available, novel genomic resources for tiny organisms such as *F. perrepae* will be more readily accessible, facilitating both fundamental genome evolutionary biology and applied sciences such as biological control programs which use eriophyoid mites for the management of invasive species.

**SIGNIFICANCE:** Eriophyid mites (Acari: Eriophyoidea) are a diverse group of miniscule mites averaging less than 0.2mm in length. The 4000 species in this clade are globally distributed and are commonly employed in biological control management of invasive species. Recent genome sequencing projects have revealed that these mites have some of the smallest genomes among animals. Here, we report on the genome assembly of the biological control eriophyoid mite *Floracarus perrepae*, which has one of the smallest genomes of any arthropod. Using this new resource, we investigate why these mites have such small genomes, focusing primarily on the loss of non-genic content such as repetitive elements and gene structure evolution. These small genomes, coupled with short generation times typical of eriophyoids, have important implications for rapid evolutionary changes, which are particularly relevant for biological control management programs.

## INTRODUCTION

The eriophyoid mites (Acari: Eriophyoidea) are a species-rich clade of more than 4000 species of minuscule arthropods (average adult length 200um, Lindquist et al. 1996) that tend to form highly specific relationships with their host plants (Skoracka et al. 2010) upon which they feed and induce the formation of galls or other plant deformities (Westphal & Manson 1996). Among mites, the eriophyoids are a morphological peculiarity, with simplified body plans specialized for phytophagy, including a reduction in the number of legs from four to two pairs, modified chelicerae, and an elongated, worm-like body plan (Figs. 1A, S1). Recent genome sequencing studies of eriophyid mites have revealed that they have some of the smallest genome sizes among animals (Greenhalgh et al. 2020; Edwards et al. 2025; Tables 1, S1). Reduction in body size and body plan, obligate relationships with their host plant, and life history characteristics such as rapid generation times (de Lillo & Skoracka 2010), could be associated with genome size reduction in this clade.

**Figure 1.**
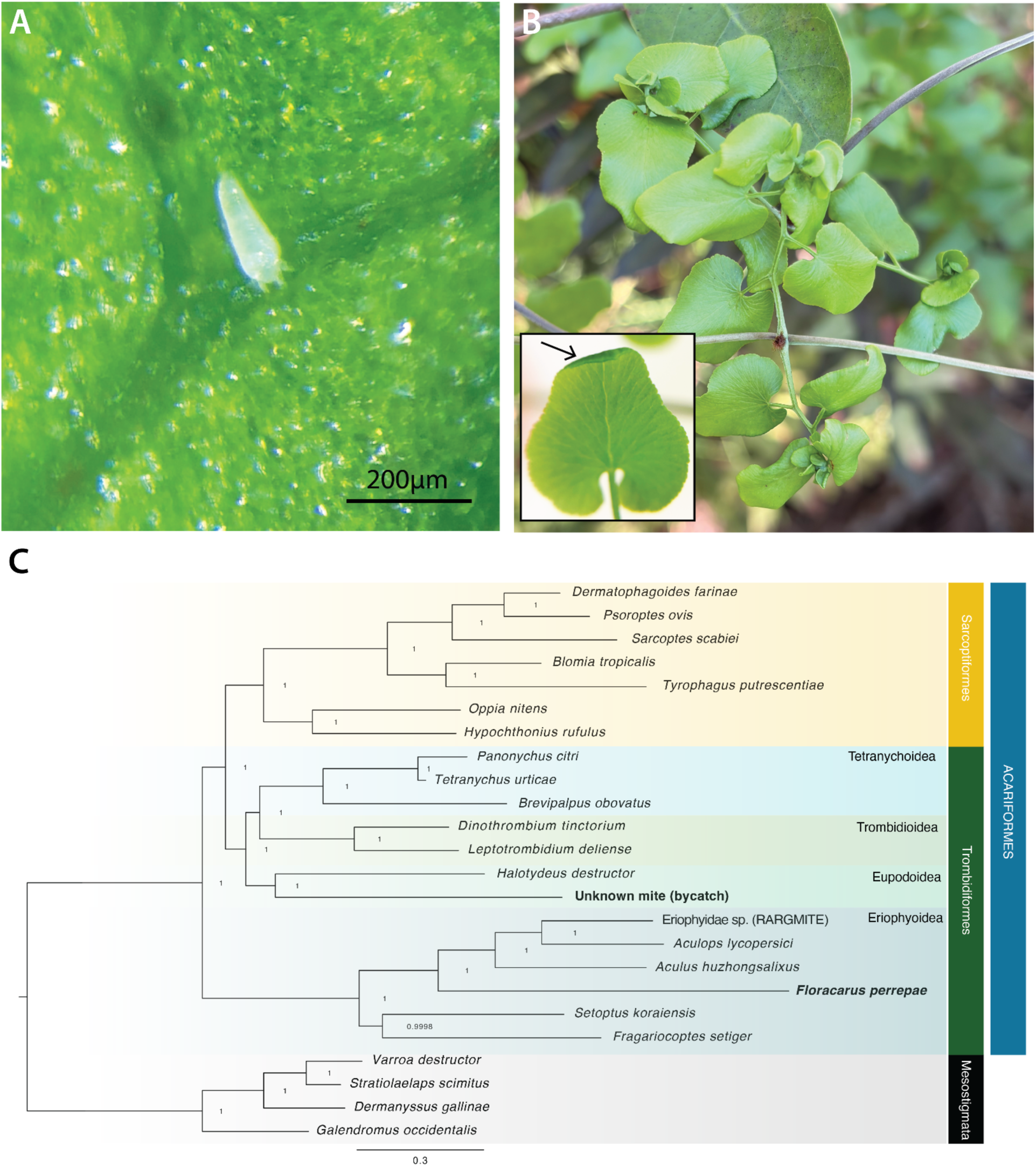
A) Photograph of *Floracarus perrepae* individual on *Lygodium microphyllum* leaflet. B) Deformities of *L. microphyllum* leaf tissue caused by *F. perrepae* feeding. Inset: arrow shows typical the galling response of *L. microphyllum* tissue caused by *F. perrepae.* C) Species tree of mite taxa generated using the multi-species coalescent, including the two genomes produced in this study in bold. Branch lengths are proportional to the number of substitutions per site and node supports are given as local posterior probabilities. The tree is rooted with taxa in the order Mesostigmata (superorder: Parasitiformes). Taxonomic groupings for superfamilies (Eriophyoidea, Eupodoidea, Trombidioidea, Tetranychoidea) are provided for the order Trombidiformes.

These characteristics also make eriophyid mites useful as candidate agents for classical and inundative biological control (Smith et al. 2010; Skoracka et al. 2010). Classical biological control involves the introduction of host-specific natural enemies (arthropod, pathogen, etc.) to elicit population-level suppression (Briese 2000; Seastedt 2015). Eriophyoid mites have been employed in classical biological control programs to varying success (e.g., Scott et al. 2008; Vidovic et al. 2016). Notably, the rapid generation times of eriophyoid mites could facilitate the rapid evolution of their ecological interactions, either beneficially (e.g., responding to plant resistance) or detrimentally (e.g., host shifts). However, studying their molecular evolution is challenging due to the difficulty in recovering DNA from such small organisms.

One such mite is *Floracarus perrepae,* a wide-spread and highly specific herbivore on the vining fern *Lygodium microphyllum* (Goolsby et al. 2003), currently used for the management this highly invasive plant (Smith et al. 2025). Associations of eriophyoid mites with ferns appear to be rare and may represent host shifts from ancestral angiosperm hosts (Gerson 1996). Individuals of *F. perrepae* are around 200μm in length (Knihinicki & Boczek 2002; Fig. 1A), have rapid generation times (9-12 days, Ozman & Goolsby 2005), reproduce year-round with a pronounced peak in spring and minor peak in fall (David et al. 2019), and are tolerant of a wide variety of climatic conditions (Ozman & Goolsby 2005; Goolsby et al. 2005; David et al. 2020). Feeding by the mite on leaflets of *L. microphyllum* induces galling, where epidermal cells become enlarged causing the margin of the leaflet to roll (Freeman et al. 2005; Fig. 1B inset). Feeding localized at the plant meristem can further induce developmental malformities (Fig. 1B). The gall acts as a protected enclosure for the mites with high humidity where they feed and reproduce, although prolonged feeding on the leaflet results in defoliation and necrosis (Freeman et al. 2005) and reduces plant climbing ability (David & Lake 2020). Dispersal to nearby plants can be accomplished by jumping (Ozman & Goolsby 2005) and alighting along wind currents (David et al. 2019). Given the hyper-specificity and stability of the interaction, *F. perrepae* has been used in control programs in the plant’s invaded range since 2008 (Boughton & Pemberton 2011), and more than 125 million mites have been released to date (Rodgers & Onisko 2026; Ralph & Klarmann 2024). While pre-release surveys found that the mite caused significant decreases in plant biomass and leaf longevity (Goolsby et al. 2004), establishment and spread has been consistent, but limited (Lake et al. 2014; Boughton & Pemberton 2011).

Here, we use a recently-developed ultra-low DNA input library preparation method to sequence and assemble a high-quality, highly-contiguous reference genome from over 200 *Floracarus perrepae* individuals, revealing one of the smallest genomes for an arthropod. We then place this new resource in the context of the existing mite genomics and explore genome downsizing in the eriophyoid mites. We anticipate that ultra-low input approaches such as ours will spur research on the evolutionary genomics of the interactions between eriophyoid mites and plants.

### Genome Assembly and Annotation of a Miniscule Mite

Galled *Lygodium microphyllum* leaf material was collected from a field site (26.09076°N, 80.41241°W) in southern Florida. Individual galls were opened and approximately 246 *Floracarus perrepae* mites were collected using a probe under a dissecting microscope. DNA was extracted from the pelleted mites using the Qiagen Qiamp Kit (Qiagen, Hilden, Germany) with an extended proteinase K digestion. Extracted DNA was sent to the Arizona Genomics Institute (AGI; Tucson, Arizona), where an ultra-low input Ampli-Fi library was constructed and sequenced on one PacBio Revio flow cell. Detailed materials and methods used in this study are described in the Supplemental Materials and Methods.

A total of 12,029,497 HiFi reads with a read N50 of 5238 bp (59.1Gb) were generated. After filtering out possible contamination from human sources and the host plant with minimap2 v2.24-r1122 (Li 2018), a total of 11,146,611 reads totaling 54.5 Gb (92.2%) were retained. The *k*-mer based estimate of genome size from KMC v3.2.4 (Kokot et al. 2017) and GenomeScope2.0 (Ranallo-Benavidez et al. 2020) using these reads was 22.5 Mb (Fig. S2), which is smaller than existing genome sequences for eriophyoid mites (Eriophyidae, 32.6-34.3 Mb, Table 1, S1). We used the metagenome assembler myloasm v0.5.1 (Shaw et al. 2026) and hifiasm v0.25.0-r726 (Cheng et al. 2021) to generate two independent assemblies from these data, which were combined with mummer v4.0.1 (Marçais et al. 2018) and quickmerge v0.3 (Chakraborty et al. 2016) (Table S2, Fig. S3). After removing haplotypic duplicates with purge_haplotigs v1.1.3 (Roach et al. 2018), the final assembly of the *Floracarus perrepae* genome was 23.13 Mb contained in 7 contigs with a contig N50 of 10.1 Mb (Table 1). The two longest contigs (10.29 Mb and 10.18 Mb) made up the majority of the assembly, with five shorter contigs comprising 11.47% (Fig. 2A). A total of 818 copies of the 5-mer AAACC (the reverse of the telomeric repeat identified by Edwards et al. 2025) were identified on one end of the longest contig assembled; no other telomeric repeats were identified at high frequency in the assembled genome. As with other eriophyoid mite genomes, the BUSCO completeness of the assembly is low (53.2% complete, Table 1), which is likely related to substantial genome downsizing (see *Genome Size Dynamism in Eriophyoid Mites*).

**Figure 2.**
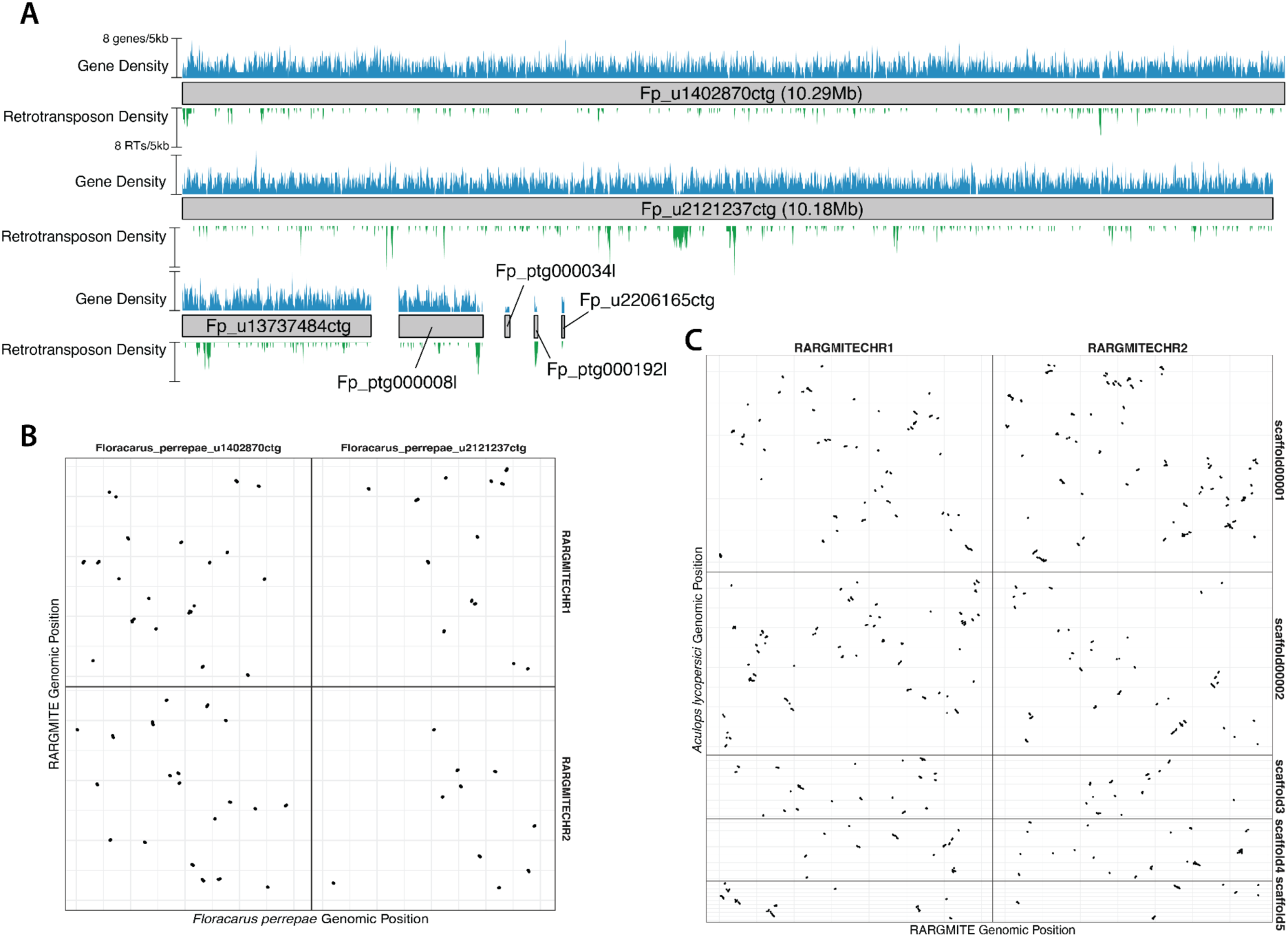
A) Contigs of *F. perrepae* reference genome painted with gene (upper, blue) and retrotransposon (lower, green) density (number of elements per 5kb). Dot plot of syntenic gene pairs between the *Floracarus perrepae* and RARGMITE (Eriophyidae sp.) genomes (B) and *Aculops lycopersici* and RARGMITE (C), where each dot represents a pair of syntenic genes; very little conserved gene order (synteny) can be observed between the eriophyoid mites.

**Table 1.** Genome assembly statistics and comparison to other eriophyoid (Eriophyoidae) mite genomes. Additional information on these and additional mite genomes used in this study can be found in Table S1.

|  | *Floracarus perrepae* | *Aculops lycopersici* | *Aculus huzhongsalixus* | Eriophyidae sp. (RARGMITE) |
| --- | --- | --- | --- | --- |
| Assembly Size (Mb) | 23,133,275 | 32,635,177 | 33,191,239 | 34,344,194 |
| Contig / Scaffold N50 | 10,182,834 | 10,499,462 | 5,931 | 17,366,404 |
| GC Content | 45.13% | 45.12% | 48.49% | 42.08% |
| BUSCO (Arachnida_ODB1 2, n=1123) | C:53.2%,F:7.7%,M:39.1% | C:59.3%,F:7.7%,M:33.0% | C:54.7%,F:11.8%,M:33.5% | C:59.5%,F:8.8%,M:31.7% |
| No. Protein-Coding Genes | 7906 | 10267 | 12119 | 8050 |
| No. Single Exon Genes (% total) | 5495 (69.5%) | 8613 (83.9%) | 8817 (72.8%) | 5257 (65.3%) |
| Reference | This study | Greenhalgh et al. 2020 | N/A | Edwards et al. 2025 |

Repetitive elements made up just 8.06% of total genomic content (Fig. 2A), most of which were simple repeats (933kb, 4.03%). Retrotransposons in the Copia (294kb, 1.27%) and Gypsy (140kb, 0.61%) families were the most abundant, but were overall uncommon when compared to other non-eriophyoid mite genomes (Table S1). Structural gene annotation with the BRAKER4 pipeline (Krall and Hoff, 2026; https://github.com/Gaius-Augustus/BRAKER4) recovered 7906 protein-coding genes, most of which were single-exon and lacking introns (5935, 75%) and 128 tRNAs, 13 rRNAs, and 93 other ncRNAs. Gene and coding sequences accounted for the majority of genome space, occupying 15.17Mb (65.59%) and 12.68Mb (54.81%), respectively. As with other eriophyoid mites, relatively few introns were annotated; 4462 introns were identified in the *F. perrepae* genome compared to an average of 4146 introns in three other Eriophyidae genomes (Table S1). On average, adjacent genes were just 1.69kb apart and 369 annotated gene pairs were overlapping (Fig. 2A). The predicted proteome had a similar completeness compared to the genome (59.10% complete BUSCOs, 65.60% complete OMark HOGs), which was close to the metrics of other eriophyoid mite genomes (Table S1).

We confirmed that the assembled genome was from *Floracarus perrepae* using a phylogenomic approach, which squarely placed the assembled genome within the Eriophyoidaea (Fig. 1C). The mitochondrial genome, assembled with MitoHiFi v3.2.1 (Uliano-Silva et al. 2023), was 16,144bp in length and contained 10 protein-coding genes, 19 tRNAs, and 2 rRNAs (Fig. S4). We verified the assembled sequence belonged to *Floracarus perrepae* using BLASTn (Camacho et al. 2009) of the assembled COX 1 sequence against the nr database on NCBI (E-value = 9e-179, 99.7-100% identity match to *F. perrepae*; Table S4).

We also recovered genome sequence of an unidentified mite as off-target bycatch. This assembly was 69.43 Mb in length (Fig. S5) composed of 101 contigs with a contig N50 of 10.90 Mb, and had a higher BUSCO completeness than the *F. perrepae* genome (84.9% complete BUSCOs). A total of 14688 protein-coding genes, 208 tRNAs, 27 rRNAs, and 169 other ncRNAs were annotated in this genome (Table S1). The predicted proteome was relatively complete (87.70% complete BUSCOs, 82.32% complete OMark HOGs), which is similar to the genome. Although the identity of this mite is unknown, best BLASTn hits of the 18s and 28s rRNAs were to superfamily Tydeoidea, which was reinforced through phylogenomic placement (Fig. 1C).

### Genome Size Dynamism in Eriophyoid Mites

Syntenic gene pairs between *Aculops lycopersici* (Greenhalgh et al. 2020)*, Floracarus perrepae*, and an unidentified eriophyoid mite (RARGMITE; Edwards et al. 2025) were identified with SynMap2 (Haug-Baltzell et al. 2017). These comparisons revealed high levels of genomic reorganization in the eriophyid mites (Fig. 2B). Only 70 syntenic blocks with an average 5.74 genes (total 402 gene pairs, 5.08% of *Floracarus* genes) were identified between *Floracarus* and RARGMITE (Fig. 2) and 157 syntenic blocks with an average of 6.39 genes (total 1004 gene pairs, 12.69% of *Floracarus* genes) were identified between *F. perrepae* and *A. lycopersici* (Fig. S6). In contrast, *A. lycopersici* and RARGMITE had 379 syntenic blocks with an average of 8.62 genes that included 3269 syntenic gene pairs (40.60% of RARGMITE genes; Fig. 2C). This is largely consistent with findings from Edwards et al. (2025), who found that gene order was largely not conserved when comparing synteny between BUSCO genes in the RARGMITE and *A. lycopersici* genomes.

To explore the evolution of genome size across the eriphyoid mites, we first used OrthoFinder v3.1.5 (Emms et al. 2026) to identify low-copy nuclear orthologs from 24 mite genomes. We then built maximum-likelihood gene trees from 302 single copy orthogroups that were present in at least 75% of taxa with IQTREE v3 (Wong et al. 2026), which were then used to construct a species tree ASTRAL IV (Zhang et al. 2025). The relationships among mites have historically been difficult to resolve (e.g., Klimov et al. 2018), especially within the Eriophyoidae, which originated sometime in the Silurian (crown age in the Permian; Zhang et al. 2024). Our goal here is not to provide a comprehensive review of these relationships, but rather to give general phylogenomic context to genome size evolution in this group. Overall, our phylogeny is similar to other mulit-gene or phylogenomic studies of Aracariformes (e.g., Edwards et al. 2025; Zhang et al. 2024; Klimov et al. 2018). We recovered monophyletic superfamilies (Eriophyoidea, Eupodoidea, Trombidioidea, and Tetranychoidea) in the Trombidiformes, but the order itself was not monophyletic (Fig. 1). Rather, Eriophyoidea was sister to a clade composed of Eupodoidea, Trombidioidea, Tetranychoidea, and Sarcoptiformes (Fig. 1). The same relationship was recovered by Edwards et al. (2025) using 1877 nuclear BUSCO genes and Zhang et al. (2024) (mitogenomes). Interestingly, Edwards et al. (2025) found that there was a high level of gene-tree conflict at the node subtending the “Trombidiformes” (Tetranycoidea + Trombidoidea) and Sarcoptiformes clade, which dates back to the boundary of the Carboniferous and Devonian (Zhang et al. 2024). Furthermore, Klimov et al. (2018) found that the placement of Eriophyoidea changed when using different datasets; their full rDNA and protein tree recovered Eriophyoidea sister to Trombidiformes + Sarcoptiformes, but their four protein dataset recovered Eriophyoidea within Trombidiformes. As genome-scale datasets become increasingly accessible for the eriophyoid mites through new sequencing approaches, resolving the placement of this species-rich and important clade should be a top priority for future studies.

We used this phylogenetic backbone to assess how genome size changed across the phylogeny and what genomic features may be associated with genome downsizing. There was strong phylogenetic signal in genome size (λ = 0.99, *P* = 2.58 x 10^-7^), with a clear and drastic reduction in genome size in the eriophyoid mites (Fig. 3A). The factors driving this downsizing in the eriophyoids remains unclear, but possible correlates could be the simplification of body plan, minute body size, obligate relationships with their host plant, and short generation times (Yang et al. 2025; Gregory & Young 2020). Decreases in repeat content are recognized as a main driver of genome size reduction (Hawkins et al. 2009; Wang et al. 2021), including in arthropods (Hancock et al. 2021). Notably, phylogenetic signal of total proportion of the genome occupied by repeats (λ = 8.13 x 10^-5^, *P* = 1), retroelements (λ = 0.17, *P* = 0.40), and DNA transposons (λ = 0.46, *P* = 0.06) were not significant, perhaps driven by lineage-specific increases in taxa such as *Dermanyssus gallinae* and *Hypochthonius rufulus*. When accounting for phylogeny, however, there was a significant association between log-transformed genome size and the proportion of the genome occupied by retroelements (PGLS, β = 0.04, adj. R^2^ = 0.38, *P* = 7.48 x10^-4^), and specifically LINEs (β = 0.04, adj. R^2^ = 0.43, *P*=2.66 x 10^-4^) but not LTRs (β = -3.9 x 10^-4^, adj. R^2^ = 0.01, *P* =0.87; Table S5). Our results show that while there is not significant phylogenetic signal in repeat content across the mites analyzed, genome size is correlated with retrotransposon content. It is possible, therefore, that suppression and excision of retroelements may have been key in genome downsizing in the eriophyoids.

**Figure 3.**
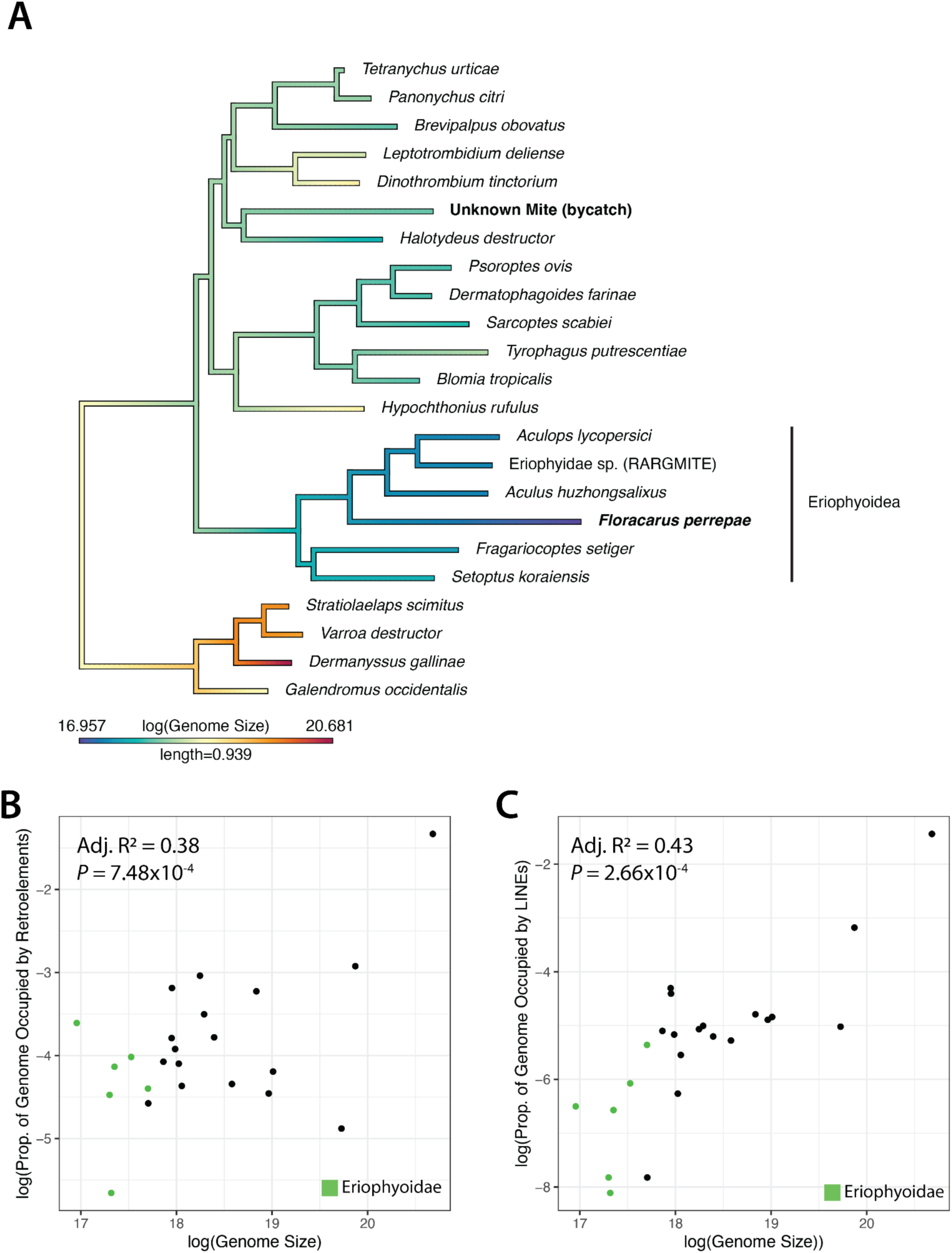
A) Phylogenetic distribution of genome size across the mite phylogeny shows distinct genome downsizing in the eriophyoid mites, and there is significant phylogenetic signal in this trait (λ = 0.99, *P* = 2.58 x 10^-7^). Genomes generated in this study are in bold. When accounting for phylogenetic relatedness, there is a significant association between genome size and the total proportion of the genome occupied by retroelements (B) and LINEs (C). The eriophyoid mites are colored green; all other mites are black.

We further investigated the evolution of gene structure evolution in light of genome downsizing. There was significant phylogenetic signal in the proportion of single-exon (i.e., intronless) genes across the phylogeny (λ = 0.99, *P* = 1.14 x 10^-6^), with a particular increase in the eriophyoid mites where up to 80% of genes lack introns in *A. lycopersici* (Greenhalgh et al. 2020), and 75% in the *F. perrepae* genome. Given the strength of this signal, we used Malin (Csurös 2008) to examine the evolution of intron site gains and losses in a phylogenetic context. A total of 47,866 intron sites were identified from 21,491 orthogroups, which were used to estimate rates of intron gain and loss at both terminal tips and internal nodes of the phylogeny. Our results suggest that there was massive intron loss along the branch subtending the eriophyoid mites (Fig. 4): the rate of loss of intron sites for the crown node of the eriophyoid mites was estimated to be 1.52 and a rate of gain of just 0.0014, for a total of 2174 lost and 147 gained intron sites at this node (Table S6). A secondary set of intron site losses also occurred at the crown node of the Eriophyidae, where an additional 394 losses and 2 gains (Table S6). Subsequent intron evolution within the Eriophyidae is relatively small compared to these changes (Fig. 4), although *F. perrepae* has the greatest number of losses (204) compared to other taxa in the Eriophyidae (Table S6). Our results highlight the utility of a phylogenetic context to understand the evolution of genome downsizing in these mites: our results suggest that evolutionary changes to gene structure likely occurred at the base of the superfamily leading to the patterns we observe in these genomes today.

**Figure 4.**
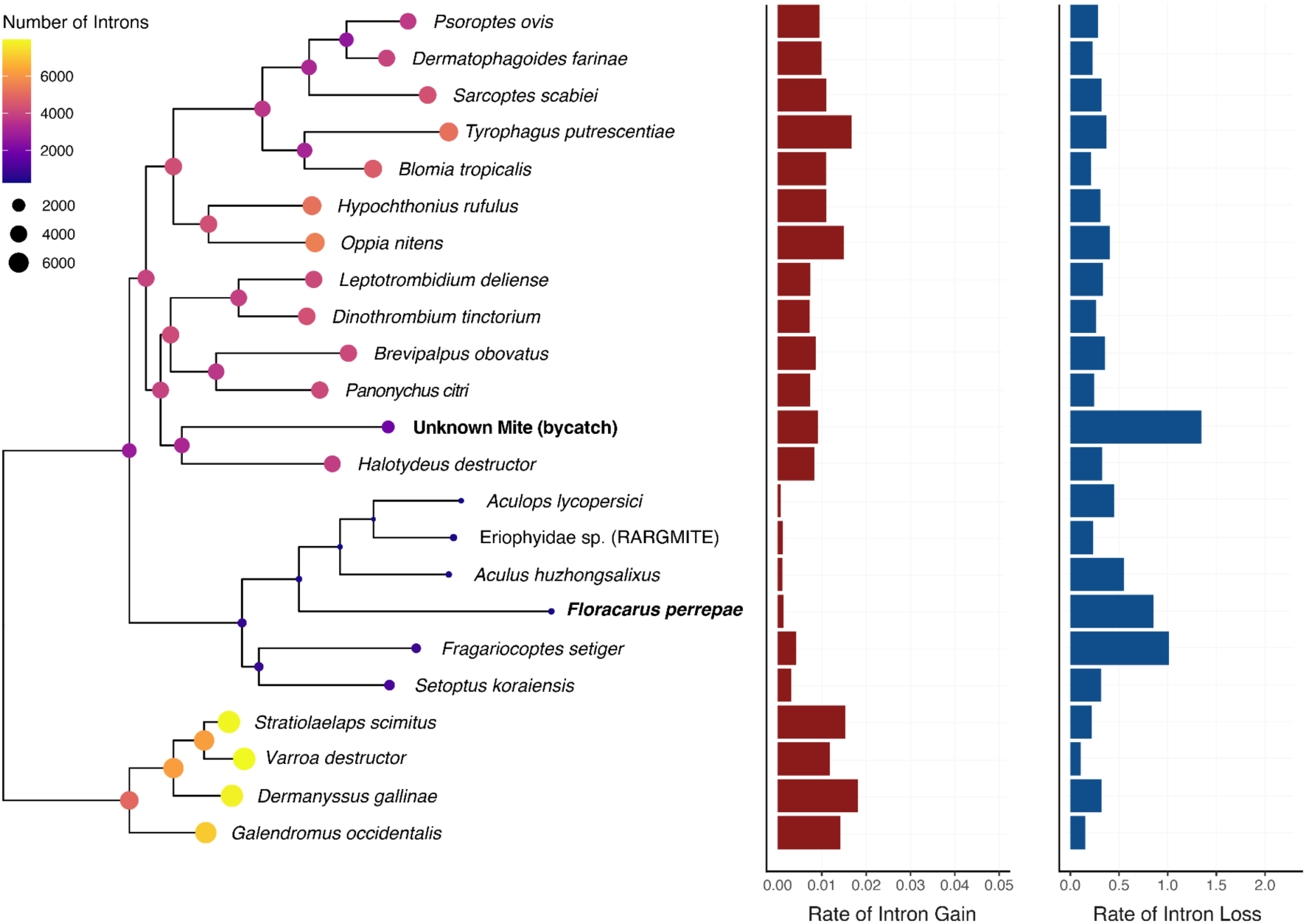
Evolution of introns across mite genomes. Circles at the nodes and tips of the phylogeny are the reconstructed number of introns present. The shape and color of the circles are scaled to this value. The rates of intron gains (left, red) and losses (blue, right) are shown to the right of the phylogeny for the tips of the phylogeny.

While we focused primarily on repeat and gene structure evolution, changes in gene family composition and loss of body-patterning genes have been associated with genome downsizing in eriophyoid mites. For example, (Greenhalgh et al. 2020) identified high rates of gene family contraction and very low rates of expansion in *A. lycopersici.* Gene families with orthologs involved in detoxification, chemoreception, and the regulation of developmental processes were rapidly contracting and missing orthologs present in other arthropods, including conserved transcription factors such as Hox genes. Notably, we also recovered strong phylogenetic signal in the proportion of the genome that is protein-coding (λ = 0.99, *P* = 7.01 x 10^-7^). Indeed, in other lineages such as nematodes (Kikuchi et al. 2017), plant-parsite species tend to have a reduction in gene number compared to non-parasitic species (Poulin & Randhawa 2015). This trend is widely observed across many parasitic species; although not necessarily in parasitic plants (Lyko & Wicke 2021). While gene loss is a common feature in parasite genomes, the expansion of gene families related to interactions with their host (e.g., in microsporidians (Nakjang et al. 2013)) and horizontal gene transfer (de la Casa-Esperón 2012) are common. Future studies should capitalize on the expanding genomic resources for the eriophyoid mites to unravel genic, as well as non-genic, evolution in this clade.

## Conclusions

Here, we generated a new highly contiguous, miniscule genome for the eriophyoid mite *Floracarus perrepae,* an active biological control agent for the invasive fern *Lygodium microphyllum*. By placing this new genome in a phylogenetic context, we revealed dynamism of eriophyoid mite genomes, including very little conservation of gene order and purging of repetitive elements and introns. Continued development of genome sequencing techniques will undoubtedly accelerate the generation of genomic resources for minute organisms such as eriophyoid mites, which will advance our understanding of genome evolution in clades where such studies were previously impossible.

## ACKNOWLEDGEMENTS

We thank Jayson Talag and Dario Copetti at the Arizona Genomics Institute for library preparation and sequencing services.

## DATA AVAILABILITY

Raw sequencing data has been deposited at NCBI under BioProject XXXXXXXXX. The genome assembly is available from NCBI under accession XXXXXXXXX, and the assembly and annotation are available at Zenodo. All code used for this project is available at XXXXXXXXX.

## SUPPLEMENTAL MATERIALS

**Supplemental Materials and Methods.**

**Supplemental Figure S1.** *Floracarus perrepae* specimen from *Lygodium microphyllum* leaf galls from field site at (26.09076°N, 80.41241°W) in Florida photographed at 1000x magnification.

**Supplemental Figure S2.** GenomeScope profile of *Floracarus perrepae* genome for *k*=21.

**Supplemental Figure S3.** Classification of contigs/components of myloasm metagenome assembly. Components 1, 3, and 4 are bycatch of an unidentified mite; Component 2 is the targeted *Floracarus perrepae* contigs. A) Assembly graph visualized in bandage. B) Blobtools results differentiate bycatch (components 1, 3, 4) from 2 (*Floracarus perrepae*) in GC content and coverage; all four components have BLAST hits to Arthopoda. C) Phylogenetic reconstruction from OrthoFinder with other mite genomes confidently places component 2 within the Eriophyoidea (sister to RARGMITE and *Aculops lycopersici*), while the bycatch assembly is most closely related to Halotydeus destructor of the included mites.

**Supplemental Figure S4.** Visualization of the annotation of the *Floracarus perrepae* mitochondrial genome.

**Supplemental Figure S5.** GenomeScope profile of mite bycatch genome for *k*=21.

**Supplemental Figure S6.** Dot plot of conserved gene order (synteny) between *Floracarus perrepae* and *Aculops lycopersici* from SynMap2.

**Supplemental Table S1.** Genome accessions and metrics of mites used in this study including *Floracarus perrepae* and the unknown bycatch.

**Supplemental Table S2.** Myloasm and hifiasm assembly results including classification of the four largest components assembled by myloasm.

**Supplemental Table S3.** Blobtools2 parsed BLAST hits for the myloasm assembly.

**Supplemental Table S4.** BLASTn results of assembled COX 1 sequence to NCBI nr database.

**Supplemental Table S5.** Results of phylogenetic signal and phylogenetically-informed regression analyses.

**Supplemental Table S6.** Results from MALIN. Highlighted nodes are those of particular interest related to the eriophyoid mites.

